# Domain-general computational integration in the Sense of Agency

**DOI:** 10.64898/2026.08.13.744643

**Authors:** Amir Harduf, Ophir Netzer, Oz Mashiah, Adam Zaidel, Roy Salomon

## Abstract

The Sense of Agency (SoA), the experience of being in control of one’s own actions, is thought to emerge from the comparison of internal sensorimotor predictions with afferent feedback. Classical comparator models treat SoA as arising from a prediction-feedback comparison. Volitional action likely engages multiple forward models that predict distinct features of an outcome, such as its timing and spatial trajectory. Whether prediction errors arising from these distinct forward models are integrated into a domain-general representation of agency, and if so by what computational logic, remains unresolved. We addressed this question using a Virtual Reality reaching task. Participants observed a virtual hand replicating their movements while we independently manipulated two sensorimotor domains: temporal delay and spatial angle deviation, in isolation and in factorial combinations. After each trial, participants made an SoA judgment. We examined whether SoA responses show computational hallmarks of integration between different features of sensorimotor prediction. Specifically, we pre-registered three computational models (Multiplicative, Minimum, and Mean) and compared their fit to per-trial responses. Across an exploratory sample (N = 16) and a pre-registered replication (N = 38), SoA declined monotonically with conflict magnitude in both domains. Critically, a Multiplicative integration rule consistently outperformed the Minimum and Mean rules. These results provide direct evidence for domain-general integration between prediction errors in SoA, governed by a multiplicative computational logic.

## Introduction

The Sense of Agency (SoA) is the subjective experience of being the initiator and controller of one’s own actions (Gallagher, 2000; Haggard, 2017). It is widely regarded as a fundamental mechanism by which individuals delineate the self from the external environment and from other agents (Zaidel and Salomon, 2023). Atypical SoA processing has been documented across a range of psychiatric and neurological conditions (Harduf et al., 2023a; Krugwasser et al., 2022a; Rossetti et al., 2026; Salomon et al., 2021; Schaefer et al., 2010), underscoring both its functional importance and its clinical relevance.

SoA has been proposed to operate at two levels: a pre-reflective, implicit *feeling* of sensorimotor control, and a higher-order, explicit metacognitive *judgment* of agency (De Vignemont and Fourneret, 2004; Legrand, 2006; Synofzik et al., 2008). Classical comparator models propose that the feeling of agency emerges when the efference copy of a motor command creates forward models to predict the sensory consequences of the action, and these predictions match the actual afferent feedback (Blakemore and Frith, 2003; Carruthers, 2012; Wolpert et al., 1995). This allows the attenuation of the sensory consequences of volitional action (Palmer et al., 2016; Shergill et al., 2012; Van Elk et al., 2014; Weiss et al., 2011) and gives rise to our sense of control. When afferent feedback deviates from these predictions, prediction errors arise. When substantial, these errors attenuate the sense of agency (Blakemore and Frith, 2003; Haggard and Chambon, 2012; Harduf et al., 2023a; Krugwasser et al., 2019; Stern et al., 2020). Consistent with this, SoA is reduced when conflict is introduced between predicted and actual action outcomes in the spatial, anatomical (Krugwasser et al., 2019; Salomon et al., 2016), or temporal (Farrer et al., 2008; Koreki et al., 2015; Limanowski et al., 2017; Stern et al., 2022; Wen et al., 2015) domains. Using Virtual Reality (VR) paradigms, for example, we have systematically introduced spatial angle deviations and temporal delays to participants’ movements and measured both explicit SoA and implicit motor responses to these perturbations (Applebaum et al., 2024).

However, a growing body of work shows that the sense of agency is also constructed from the integration and weighted combination of multiple cues, including efferent signals, proprioceptive and visual feedback, contextual information, and prior beliefs about action outcomes (Moore et al., 2009; Moore and Fletcher, 2012; Synofzik et al., 2013, 2008). A comprehensive account of SoA, therefore, requires characterizing not only individual prediction-feedback comparisons but also how multiple agency-relevant signals interact (**Fig. 1a**). This raises a question that remains largely unresolved: Is there a domain-general representation of SoA that integrates across distinct forward models? The sensory predictions arising from volitional action span multiple domains and require different computations to compare efferent and afferent signals. Moving a hand, for example, engages forward models predicting *which* hand moved (anatomical), *when* it moved (temporal), and *where* it moved (spatial). These models draw on different signals and can, in principle, generate independent prediction errors: one may move the correct hand at the correct time, yet along a perturbed spatial trajectory. Whether prediction errors across these distinct forward models are integrated, and if so, by what computational logic (**Fig. 1d**), remains an open question. Several studies provide evidence for a domain-general representation. Signal-detection analyses have shown that both sensitivity (d′) and bias for detecting action perturbations are strongly correlated across anatomical, temporal, and spatial domains (Krugwasser et al., 2019; Stern et al., 2020), suggestive of a shared domain-general computation. In parallel, recent neuroimaging work using fMRI (Harduf et al., 2023b) and EEG (van der Goot et al., 2026) has supported the two-step model, identifying dissociable neural signatures of low-level prediction-error processing and higher-level agency judgments. What is still missing is a direct demonstration of *integrations* between prediction errors arising from distinct forward models and a formal characterization of the computational logic that governs their integration.

**Figure 1.**
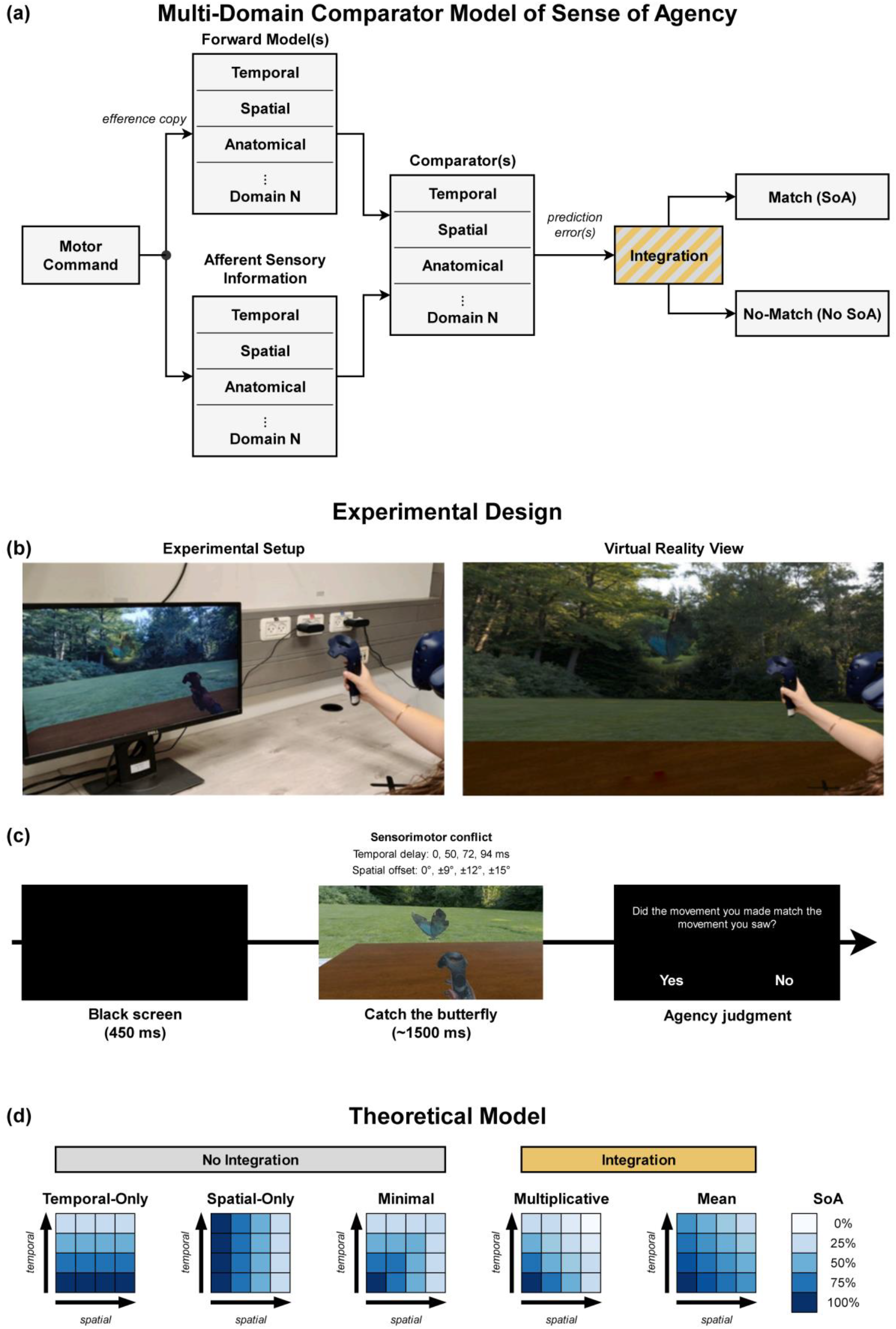
Multi-domain comparator model, experimental design, and candidate models. **(a)** Multi-domain comparator model of the sense of agency. Forward model(s) predict the sensory feedback of a motor command across domains (temporal, spatial, anatomical, etc.); domain-specific comparator(s) compute the prediction error against the actual afferent feedback. These errors feed a putative integration mechanism (yellow/grey) that yields Match (agency) or No-Match (no agency), the step formalized by the candidate models in panel d (yellow = integration, grey = non-integration). **(b)** Experimental Design. Left: a participant performing the task in the apparatus. Right: an illustration of the participant inside the virtual-reality environment. **(c)** Trial procedure: a black screen, then a virtual “catch the butterfly” reaching task, then an agency judgment. Sensorimotor conflict was manipulated along two domains: temporal delay and spatial offset. (**d**) Candidate model predictions. Predicted self-attribution across the temporal (rows) × spatial (columns) alteration grid for five models, grouped into non-integration models in gray (Temporal-Only, Spatial-Only, Minimal) and integration models in yellow (Multiplicative, Mean). Cell shading in the blue palette indicates predicted SoA.

Here, we directly test whether prediction errors arising from distinct forward models are integrated into a domain-general representation of agency, and we adjudicate between competing computational accounts of how this integration is implemented. Focusing on the temporal and spatial domains, we used an established VR reaching task (Applebaum et al., 2024; Stern et al., 2024) in which participants observed a virtual hand replicating their movements, while we independently manipulated temporal delay and spatial angle deviation, both in isolation and in factorial combination (**Fig. 1b, c**). Participants made a binary self-attribution judgment on each trial. We hypothesized that (H1) SoA would decline monotonically with conflict magnitude in each modality, and (H2) that a model integrating temporal and spatial prediction errors within a trial would outperform non-integration models. To test H2, we fit three pre-registered computational models: Multiplicative, Minimum, and Mean, to per-trial responses and compared them using BICs (**Fig. 1d**). Across both an exploratory sample (N = 16) and a pre-registered replication (N = 38), a multiplicative integration rule best accounted for self-attribution judgments, outperforming the non-integration Minimum model, which uses both signals but lets the larger prediction error alone determine agency in a winner-take-all fashion without combining them, and the Mean model, an averaging-based integration rule. These results provide direct evidence for domain-general integration of multimodal prediction errors in the sense of agency, and suggest that this integration follows a multiplicative rather than averaging or winner-take-all computational logic.

## Methods

### Participants

Two cohorts participated. In experiment 1, 16 neurotypical adults were recruited. For the pre-registered replication (preregistration: https://aspredicted.org/kq779q.pdf), we targeted N = 34 based on power analyses (alpha=.05, power=.80) using effect sizes from the exploratory phase: the required N was 32.57 for the temporal Friedman test (Kendall’s W = 0.704), 24.25 for the spatial Friedman test (W = 0.821), and 14.7 for the primary model comparison (Cohen’s d = 1.07). N = 38 was selected to satisfy all three estimates and enable counterbalancing. Data collection continued until 38 valid participants were obtained; all 38 completed the study and met the inclusion criteria. All participants were right-handed and had normal or corrected-to-normal vision. Prior to participation, informed consent was obtained, and individuals were compensated either with 50 NIS per hour or with course credits. All procedures were performed in compliance with the institutional guidelines of the Gonda Multidisciplinary Brain Research Center at Bar-Ilan University (BIU) and the University of Haifa. The study was reviewed and approved by the respective institutional ethics committees (BIU approval number: ISU202107003; University of Haifa approval number: 283/23).

### Experimental Setup

To create a highly immersive and realistic virtual environment using in-house software developed in Unity (2018.3.2), participants were seated at a physical table and equipped with a Vive head-mounted display (90Hz refresh rate; **Fig. 1b**). Two base stations tracked participants’ hand motions to generate the environment. Participants held a Vive controller in their right hand, initiating and concluding each trial’s movement from a designated black ‘x’ marker on the table. The controller’s trackpad was employed as an input device to record their answers. To maintain immersion, the left hand remained hidden beneath the physical table, as only the right avatar hand was displayed in VR. The physical workspace was precisely aligned with its virtual counterpart, matching the table, ‘x’ marker, and controller, to ensure congruent visual and tactile feedback. From this calibrated starting position, participants utilized their virtual hand to catch a butterfly hovering at a designated distance above the table.

### Procedure

Participants performed a VR reaching task toward a virtual butterfly target. Each trial (**Fig. 1c**) began with a black screen (600ms) followed by the virtual environment. After 200ms, the butterfly appeared, ~60 cm from the hand’s starting position. As participants reached toward the butterfly, it disappeared once the hand had moved ~20 cm from its starting position, to prevent it from being used as a feedback cue. Throughout the movement, participants observed a virtual hand replicating their trajectory, with one or both types of sensory-motor conflict introduced per trial. Following the movement, participants answered a binary forced-choice question presented in Hebrew (“Did the movement you made match the movement you saw?”; Yes/No).

The task comprised three sequential blocks. The order of the first two blocks was counterbalanced across participants. Block 1 (Temporal-only, 72 trials) held spatial conflict at 0° while temporal delay was varied across four levels (0, 50, 72, 94 ms). Block 2 (Spatial-only, 72 trials) held temporal delay at 0 ms while spatial angle deviation was varied across four levels (0°, +/-9°, +/-12°, +/-15°). Block 3 (Combined, 234 trials) presented all 16 combinations of the four temporal and four spatial levels, with baseline trials (0 ms, 0°) constituting approximately 25% of the block. The virtual target was presented at 3 counterbalanced positions along the vertical axis (bottom, middle, top), all at the same horizontal position. Post-task, participants completed the IPASE (Inventory of Psychotic-Like Anomalous Self-Experiences), PQ-B (Prodromal Questionnaire-Brief), OCI-R (Obsessive-Compulsive Inventory-Revised), and an in-house demographics questionnaire.

Participants completed a training phase of 20 trials with up to three attempts. To be included in the final analysis, they were required to meet level-specific accuracy criteria across the four difficulty levels. For participants who reached a third attempt, the criterion was waived at the highest difficulty level and raised at an intermediate one, retaining those who could not detect the subtlest perturbations while still maintaining reliable discrimination elsewhere. Participants were excluded if they failed to pass the training or did not complete all experimental blocks (for a detailed procedure, see the preregistration: https://aspredicted.org/kq779q.pdf).

### Computational Modeling

Three pre-registered models were fit to each participant’s Combined block data using maximum likelihood estimation with the BADS (Bayesian Adaptive Direct Search) optimizer (Acerbi and Ma, 2017; four random restarts; negative log-likelihood objective). All models used monotonicity-constrained parameters. Each model combined the fitted per-level parameters (T_level_, S_level_ ∈[0,1]; the temporal and spatial parameters at the level of conflict presented on that trial) into a predicted self-attribution probability; Multiplicative: P = T_level_ × S_level_, Minimum: P = min(T_level_, S_level_), and Mean: P = (T_level_ + S_level_)/2. Each had eight free parameters (four T_level_, four S_level_). Model fit was evaluated using the Bayesian Information Criterion: BIC *= −2*log(L) + log(N)*k, where L is the maximized likelihood, k is the number of free parameters, and N is the number of trials.

### Statistical Analysis

Normality was assessed with Shapiro-Wilk tests; parametric tests were used when normality held and non-parametric alternatives otherwise. To test whether self-attribution rates differed across alteration levels, Friedman tests were applied to each participant’s mean self-attribution rate across the four alteration levels, separately for the temporal and spatial blocks, with Kendall’s W as the effect size. Significant omnibus tests were followed by post hoc Wilcoxon signed-rank tests for all pairwise level comparisons, with Benjamini-Hochberg FDR correction. As a preliminary test of whether the effect of one conflict depended on the level of the other, a mixed-effects logistic regression (GLMM; soa_rating ~ delay x angle + [1|Subject]) was fit to the Combined block. Significant main effects would indicate that participants used both domains, ruling out theoretical unimodal accounts in which judgments are driven by a single domain (**Fig. 1d**), and a significant interaction term would indicate that the two domains were not processed independently, motivating the model comparison that follows. Full results are reported in the Supplementary Information. To test the nature of these interactions and whether these domains were combined via a model that integrates both domains within a trial (e.g., Multiplicative, Mean), yielding better predictions than models that do not integrate both domains (e.g., Minimum), per-participant ΔBIC was computed for each model relative to the others. Comparisons used Wilcoxon signed-rank tests (non-normal) against mu = 0. The primary pre-registered test was Multiplicative vs. Minimum; secondary tests compared Multiplicative vs. Mean, and Mean vs. Minimum. All tests used alpha = .05, two-tailed, with FDR correction, based on three comparisons.

## Results

### Unimodal Sense of Agency Declines with Temporal Delay and Spatial Offset

Self-attribution rates decreased monotonically as sensory-motor conflict increased in both domains, with self-attribution falling by more than half from baseline to the largest alteration (**Fig. 2a, b)**. In experiment 1 (N = 16), Friedman tests confirmed significant differences across alteration levels for both the temporal (χ^2^ (3) = 33.84, p = 2.1×10^−7^, W = 0.705) and spatial (χ^2^ (3) = 39.44, p = 1.4×10^−8^, W = 0.822) conditions, with all pairwise comparisons between levels significant (all p < .05, FDR-corrected). The replication sample (N = 38) yielded similar results. Self-attribution declined significantly across temporal (χ^2^ (3) = 78.07, p = 7.9×10^−17^, W = 0.685) and spatial (χ^2^ (3) = 71.95, p = 1.6×10^−15^, W = 0.631) conditions, with all pairwise comparisons between levels significant (all p < .01, FDR-corrected; **Fig. 2c, d**).

**Figure 2.**
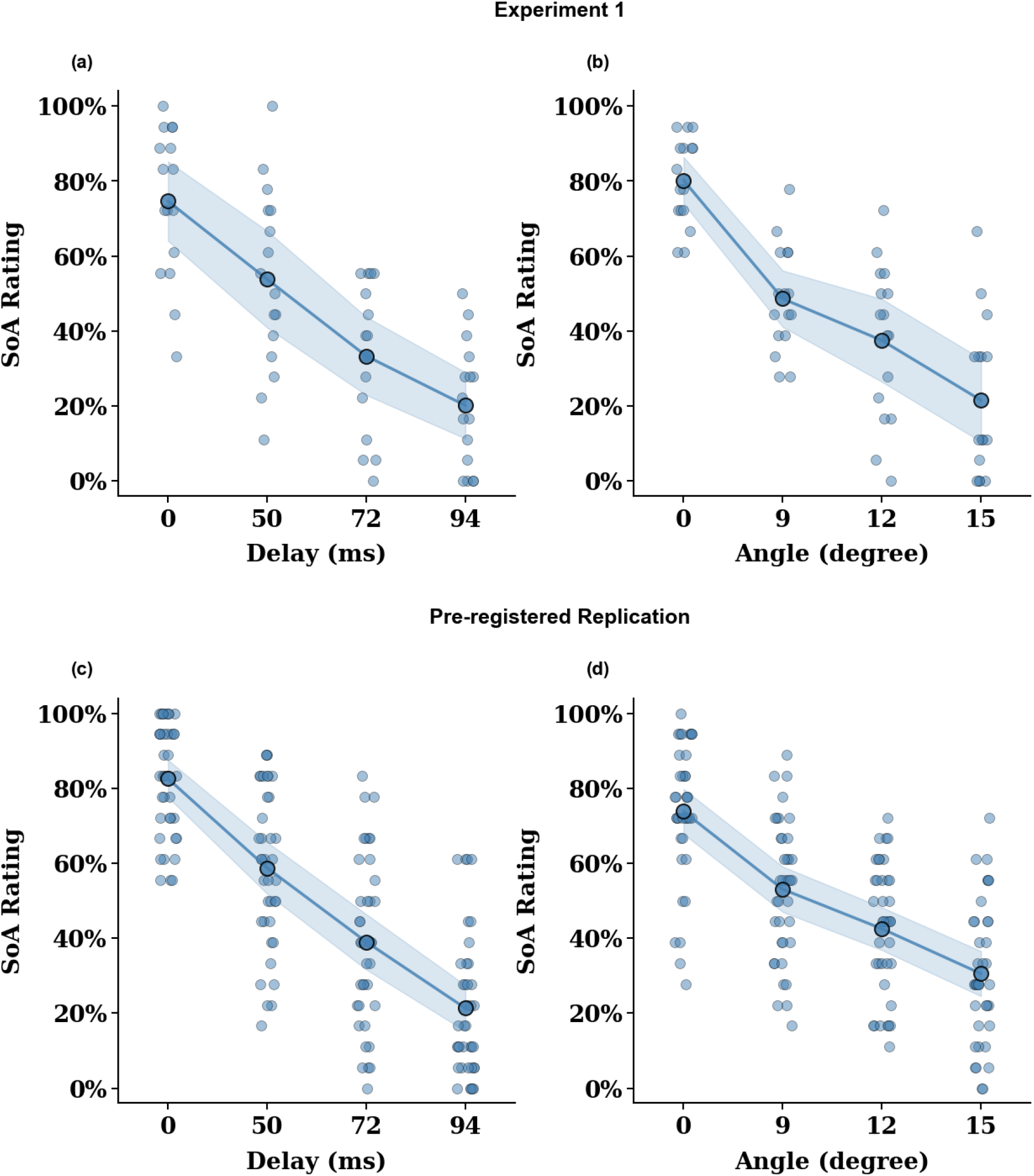
Self-attribution across unimodal alteration levels. Self-attribution rate (percentage of “Yes” agency responses) at each level of temporal delay (Temporal-only block) and spatial offset (Spatial-only block). Blue markers represent the group means at each level (connected across levels), the shaded band the 95% confidence interval, and grey dots individual participants. Top row: Experiment 1 (N = 16); bottom row: pre-registered replication (N = 38). **(a, c)** Temporal block; **(b, d)** Spatial block.

### Multiplicative Integration Model Outperforms Non-Integration and Unimodal Models

In both samples, self-attribution declined with temporal and spatial conflict, and the effect of each depended on the level of the other (GLMM; all main effects and the interaction p < .05; **Supplementary Results**). Both conflicts affected self-attribution, ruling out a unimodal account (**Fig. 1d**), while the interaction indicates that the two domains were not processed independently. We therefore compared candidate models to characterize how they were combined. To test whether the two domains were integrated within a trial, we compared two integration models (Multiplicative, Mean) against a non-integration baseline (Minimum) using per-participant ΔBIC. The primary pre-registered comparison, Multiplicative vs. Minimum, favored Multiplicative in both samples (Experiment 1: p = .004, FDR-corrected, d = −1.07; Replication: p = .041, FDR-corrected, d = −0.42; **Fig. 3** and **Supplementary Tables 1 and 2**). Multiplicative also outperformed the averaging (Mean) model in both samples (Experiment 1: p = .014, FDR-corrected, d = −0.54; Replication: p = .0006, FDR-corrected, d = −0.59). In contrast, the Mean model did not outperform the Minimum baseline (Experiment 1: p = .744, FDR-corrected, ns; Replication: p = .032, FDR-corrected, favoring Minimum), indicating that simple averaging does not capture the integration. Thus, across both samples, only the multiplicative rule consistently outperformed the non-integration and alternative-integration models, thereby identifying the multiplicative combination as the integration rule.

**Figure 3.**
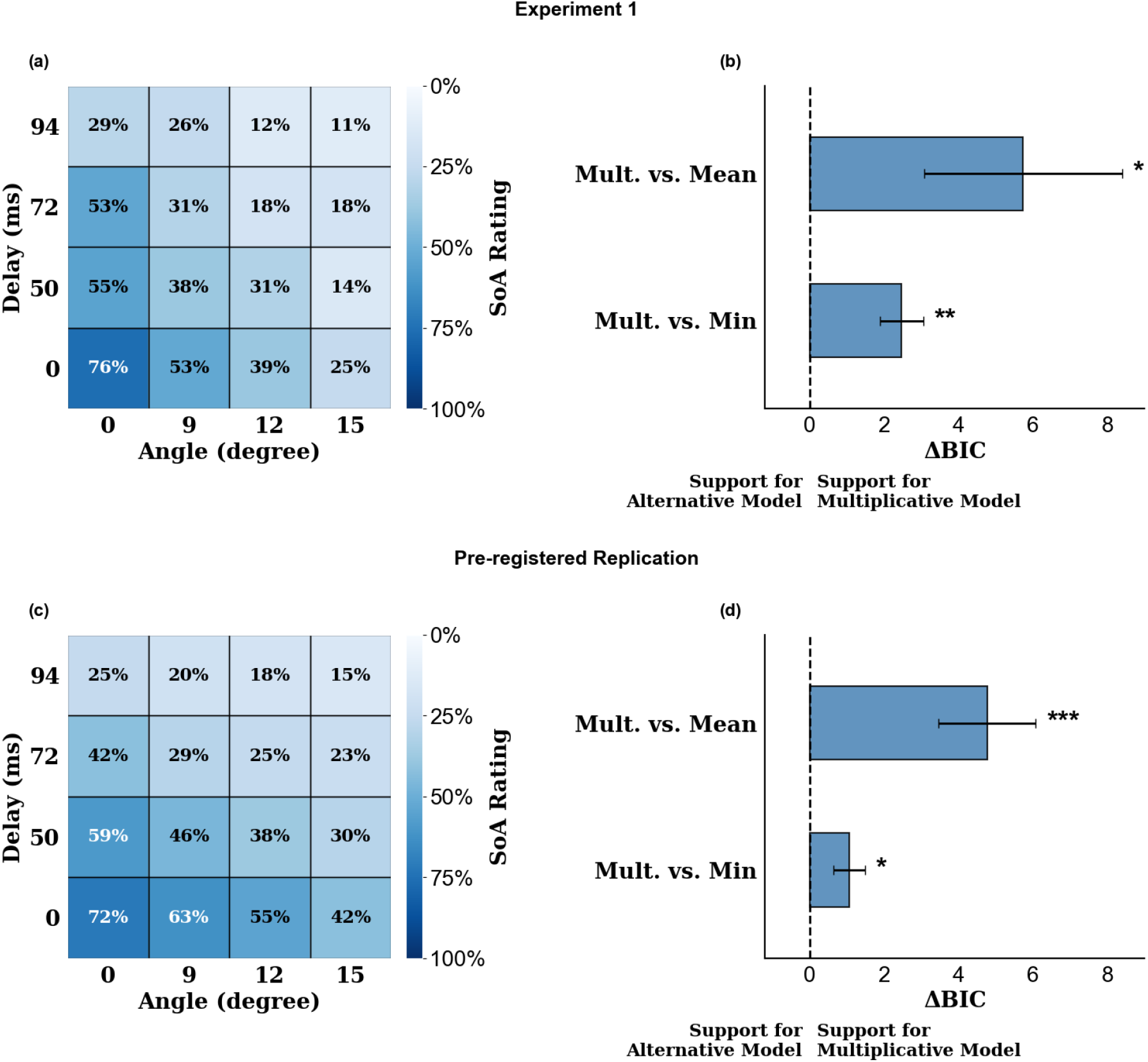
Combined-block self-attribution and model comparison. **(a, c)** Mean self-attribution rate across the temporal delay × spatial offset grid of the combined block; cell shading indicates self-attribution (color bar). **(b, d)** Model comparison by ΔBIC: bars show the mean per-participant ΔBIC (± SEM) of the Multiplicative model relative to the Minimum and Mean models (bar labels: Mult. = Multiplicative, Min = Minimum); positive values indicate better fit (lower BIC) for the Multiplicative model. Asterisks denote FDR-corrected significance (* p < .05, ** p < .01, *** p < .001). Top row: Experiment 1 (N = 16); bottom row: pre-registered replication (N = 38).

## Discussion

The present study tested whether prediction errors arising from distinct sensorimotor conflicts are integrated into a domain-general representation of agency, and if so, by what computational logic. Three findings emerged. First, as expected, SoA declined monotonically with conflict in both the temporal and spatial domains, replicating prior single-modality work within a factorial design. Second, a multiplicative integration rule fit per-trial judgments better than unimodal, winner-take-all, and averaging-based alternatives, across both an exploratory and a pre-registered replication sample. Together, these results indicate that the comparator framework, while correct in positing prediction-feedback comparison, is incomplete: prediction errors generated by separate forward models are not evaluated independently, but jointly determine agency through a conjunctive, multiplicative rule (**Fig. 1a**).

First, as expected, SoA judgments declined monotonically with conflict magnitude in both the temporal and spatial domains, replicating prior single-modality work (Farrer et al., 2008; Kannape et al., 2010; Krugwasser et al., 2019; Stern et al., 2020) within a factorial design. The effect was large and replicated across samples, confirming that participants can give stable explicit agency judgments corresponding to graded levels of visuomotor conflict.

The model comparison adjudicates between two theoretically distinct possibilities. On the one hand, it is possible that different aspects of sensorimotor error are processed separately and are not integrated. Alternatively, if some form of integration between the error signals is present, this would constitute strong evidence of supramodal processing of SoA. Indeed, in both experiments an integration model outperformed the non-integration model. The Minimum model provided a poorer fit to the data, indicating that SoA was not determined solely by the signal with the smallest prediction error. Instead, temporal and spatial prediction errors jointly shaped agency judgments, supporting genuine integration rather than winner-take-all selection, and suggesting a supramodal cognitive integration process.

When examining the computational logic of the integration process, we compared two competing accounts: a Mean model and a Multiplicative model. Both assume that temporal and spatial prediction errors are integrated into a common agency estimate, but they differ fundamentally in how this integration is computed. The Mean model assumes compensatory, additive integration, whereby a strong match in one modality can offset a mismatch in the other. In contrast, the Multiplicative model implements conjunctive integration, in which both prediction signals jointly determine the final sense of agency, such that a mismatch in either modality substantially reduces the agency estimate. Our results consistently favored the Multiplicative model across both the exploratory and replication samples, demonstrating that prediction errors are not simply averaged but combined through a conjunctive computational rule. This suggests that the sense of agency depends on the joint consistency of multiple forward-model predictions rather than on the average strength of individual prediction signals.

These findings argue for extending the comparator model for SoA (**Fig. 1a**). The classical formulation is centered on the conceptual idea that a single prediction-feedback comparison generates a single error signal (Frith, 2012; Gentsch et al., 2012; Wolpert et al., 1995). Our results indicate that volitional action engages multiple forward models in parallel, each producing its own prediction error, and that SoA emerges from a downstream integration stage that combines these errors multiplicatively. This architecture is consistent with prior evidence for domain-general computation, including shared sensitivity and bias across domains (Krugwasser et al., 2022b, 2019; Stern et al., 2022) and dissociable neural signatures of prediction-error processing and agency judgment (Harduf et al., 2023b; van der Goot et al., 2026). The present work provides a direct behavioral demonstration of the integration itself, and a formal characterization of its computational rule.

Four limitations bound the present conclusions. First, only two domains were manipulated; whether anatomical or prediction errors from other (e.g., auditory or somatosensory) modalities are integrated by the same rule remains an open question. Second, agency was indexed by explicit binary judgments; implicit measures such as sensory attenuation (Bansal et al., 2018; Kilteni et al., 2020; Kilteni and Ehrsson, 2017; Weiss et al., 2011) or hand movements (Applebaum et al., 2024) may follow different integration logic and warrant direct comparison. Third, the conflict ranges were calibrated near the detection threshold to maximize sensitivity; generalization to the suprathreshold perturbations of everyday action is not guaranteed.

In conclusion, our work reveals that prediction errors arising from temporally and spatially distinct forward models are integrated into a domain-general representation of agency, and this integration follows a multiplicative rather than additive or winner-take-all rule. These findings extend the comparator model from a single prediction-feedback comparison to a multi-stage architecture in which independent forward-model errors are conjunctively combined to yield the agency judgment.

## Acknowledgment

This work was supported by an ERC STG (UNREAL-949010) to R.S

## Notes

### Competing Interest Statement

The authors have declared no competing interest.

