## Supplementary Information for "Domain-general computational integration in the Sense of Agency"

### Supplementary Results: Mixed-effects logistic regression (GLMM)

A mixed-effects logistic regression (soa_rating ~ delay × angle + [1|Subject]) was fit to the Combined block in each sample. Both domains produced strong main effects. In Experiment 1 ($N = 16$), self-attribution declined with both temporal ($\beta = -0.021$, $z = -13.04$, $p < .001$, $OR = 0.98$) and spatial ($\beta = -0.154$, $z = -15.77$, $p < .001$, $OR = 0.86$) conflict, baseline ≈ 79% (intercept $\beta = 1.33$, $z = 7.20$, $p < .001$), with a small but significant temporal × spatial interaction ($\beta = 0.0007$, $z = 3.74$, $p < .001$). The replication ($N = 38$) reproduced both main effects (temporal: $\beta = -0.021$, $z = -20.45$, $p < .001$, $OR = 0.98$; spatial: $\beta = -0.076$, $z = -13.03$, $p < .001$, $OR = 0.93$; baseline ≈ 75%, intercept $\beta = 1.09$, $z = 10.38$, $p < .001$) and the interaction ($\beta = 0.0002$, $z = 2.01$, $p = .044$). The significant main effects in both samples indicate that participants used both domains, ruling out a theoretical unimodal account (**Fig. 1d**). The significant interaction indicates that the effect of each conflict depended on the level of the other; its form is characterized by the model comparison reported in the main text.

### Supplementary Table 1: Experiment 1 - Pre-registered ΔBIC comparisons

| Comparison | ΔBIC | SEM | d | p_raw | p_FDR | Significance |
|---|---|---|---|---|---|---|
| **Multiplicative vs. Minimum** | 2.47 | 0.58 | 1.07 | 0.0013 | 0.004 | ** |
| **Multiplicative vs. Mean** | 5.74 | 2.66 | 0.54 | 0.009 | 0.014 | * |
| **Mean vs. Minimum** | -3.27 | 2.56 | -0.32 | 0.744 | 0.744 | ns |

**Supplementary Table 1: Experiment 1 - Pre-registered model comparisons.** Per-participant ΔBIC between integration models (positive favors the Multiplicative model). Wilcoxon signed-rank; FDR-BH across 3 comparisons.

### Supplementary Table 2: Replication - Pre-registered ΔBIC comparisons.

| Comparison | ΔBIC | SEM | d | p_raw | p_FDR | Significance |
|---|---|---|---|---|---|---|
| **Multiplicative vs. Minimum** | 1.06 | 0.42 | 0.42 | 0.041 | 0.041 | * |
| **Multiplicative vs. Mean** | 4.76 | 1.31 | 0.59 | 0.0002 | 0.0006 | *** |
| **Mean vs. Minimum** | -3.7 | 1.42 | -0.42 | 0.022 | 0.032 | * |

**Supplementary Table 2: Replication - Pre-registered model comparisons.** Per-participant ΔBIC between integration models (positive favors the Multiplicative model). Wilcoxon signed-rank; FDR-BH across 3 comparisons.
